# Spatial Omics Driven Crossmodal Pretraining Applied to Graph-based Deep Learning for Cancer Pathology Analysis

**DOI:** 10.1101/2023.07.30.551187

**Authors:** Zarif Azher, Michael Fatemi, Yunrui Lu, Gokul Srinivasan, Alos Diallo, Brock Christensen, Lucas Salas, Fred Kolling, Laurent Perreard, Scott Palisoul, Louis Vaickus, Joshua Levy

## Abstract

Graph-based deep learning has shown great promise in cancer histopathology image analysis by contextualizing complex morphology and structure across whole slide images to make high quality downstream outcome predictions (ex: prognostication). These methods rely on informative representations (i.e., embeddings) of image patches comprising larger slides, which are used as node attributes in slide graphs. Spatial omics data, including spatial transcriptomics, is a novel paradigm offering a wealth of detailed information. Pairing this data with corresponding histological imaging localized at 50-micron resolution, may facilitate the development of algorithms which better appreciate the morphological and molecular underpinnings of carcinogenesis. Here, we explore the utility of leveraging spatial transcriptomics data with a contrastive crossmodal pretraining mechanism to generate deep learning models that can extract molecular and histological information for graph-based learning tasks. Performance on cancer staging, lymph node metastasis prediction, survival prediction, and tissue clustering analyses indicate that the proposed methods bring improvement to graph based deep learning models for histopathological slides compared to leveraging histological information from existing schemes, demonstrating the promise of mining spatial omics data to enhance deep learning for pathology workflows.

## 1. Introduction

### 1.1. Deep Learning for Pathology

In recent years, countless studies have demonstrated the potential for deep learning algorithms to solve challenging biomedical tasks, thereby improving risk stratification and alleviating the potential for clinical burnout by making tedious and unreliable tasks faster and more quantitative, potentially leading to improved patient health outcomes ^1^. These algorithms are formulated on computational heuristics – specifically, machine learning -- which can make sense of many complex data types through the dynamic derivation of relevant patterns and features ^2–4^. Analysis of pathology data, including whole slide imaging (WSI) – microscopic images of patient tissue – is an emerging application in this space, as WSIs are routinely collected and used for patient monitoring, diagnosis, and prognostication. Existing works have shown that specially designed deep learning algorithms, inspired by processes of the central nervous system, may be able to automate or assist in these tasks ^5^. Most deep neural networks study small micromorphological changes given the enormity of these gigapixel images. Graph convolutional networks (GCNs), however, are a promising method in this domain, as they can effectively model macro and micro architectural features present across WSI in a human-interpretable manner ^6–8^. Generally, these methods split WSI into patches (i.e., more manageable subimages), extract numeric representations (i.e., “embeddings”) from each patch using a predetermined algorithm, and construct a graph where the nodes are given patch embeddings and edges are formed based on spatial adjacency.

The optimal algorithm used to extract node features is an area of ongoing research, though many works presently use a ResNet convolutional neural network (CNN) pretrained on the ImageNet database ^9^ for this task. It has become increasingly common to additionally train these CNNs on various image tasks orthogonal to the task at hand to prepopulate an information registry of features which will ultimately improve predictive performance in other settings; these techniques are known as pretraining. Recently, self-supervised techniques have emerged as promising pretraining methodologies, where images are compared form several different vantage points without being explicitly labeled ^10^. Cross-modal pretraining has recently been highlighted as a common self-supervised method by leveraging complementary “paired” information across multiple input data types (e.g., images and text) which can improve the representation of all involved modalities. Here, we investigate the utility of using spatial omics data, which is paired at 50-micron resolution to the histological information, to pretrain an encoder model for these patches, to demonstrate the power of leveraging spatial omics for deep learning-based pathology methods which are particularly suited for analysis using graph neural networks (GNNs).

### 1.2. Spatial Omics

Omics data – such as gene expression quantification and DNA methylation – have traditionally been collected on a bulk scale where measurements are taken across an entire sample or tissue section. Recent advancements in technology have allowed for collection on a more granular scale, such as the single cell level, or across specific spots/regions in a slide sample ^11^. Prior studies have demonstrated that deep learning through specialized architectures like GCNs can mine spatial omics data to build a more comprehensive understanding of spatial cellular heterogeneity, especially as it pertains to how the tumor microenvironment can facilitate/inhibit further disease progression ^12,13^. Notably, this type of data is not yet commonly available at large scale due to the prohibitive cost of these assays as well as batch effects and selection of limited slide area, meaning that methods which can learn from spatial omics data and effectively transfer this knowledge to improve other tasks may be valuable. Zeng et al ^13^ previously developed a model which utilized contrastive learning to mine a shared representation between image patches and corresponding spatial transcriptomics; however, their investigation centered on driving improved understanding on gene domains, rather than attempting to leverage the method to enhance downstream clinical outcome modeling in situations where only WSI – and no ST data – is available.

### 1.3. Contributions

We hypothesize that additional biological information can be learnt from spatially resolved transcriptomics data that may prove relevant for enhancing prediction models across a range of histological analyses. Existing works applying GCNs for WSI analysis have not yet leveraged spatial omics data to enhance modeling across orthogonal tasks. In part, this is because the quality of histological slides for spatially co-registered omics data has been limited as the standard Visium spatial transcriptomics (ST) workflow featured manual staining and low-resolution imaging– this information does not readily transfer to prediction models on higher resolution histological slides. Now, with the development of assays such as the CytAssist which permit the use sophisticated laboratory processing (i.e., autostaining and 40X imaging prior to Visium profiling), the quality of slides has remarkably increased and allows for training image models that may more readily transfer to related domains. Here, we assess the ability of spatially resolved omics data to enhance predictions on a range of different histological assessment tasks by presenting an initial evaluation of a crossmodal pretraining mechanism using matched WSI and spatial omics measurements as means to encode biological information within WSI graphs. We compare this method against other common pretraining schemes on downstream predictive analyses (staging, lymph node metastasis, survival prognostication) of WSI, as well as explore generated image patch embeddings. Accurate methods for these downstream predictive tasks may enable more personalized patient treatments. In this study, we expect developed models which can mine for spatial molecular information to outperform the compared approaches on these tasks. We aim to demonstrate the potential benefits of utilizing spatial omics – spatial transcriptomics, in particular – methods to enhance deep learning-driven pathology analysis.

## 2. Methods

### 2.1. Data Collection and Preprocessing

Visium spatial transcriptomics data matched with WSI at 40x resolution was collected from 8 colorectal cancer patients from the Dartmouth Hitchcock Medical Center, to serve as a training dataset for the crossmodal patch embedding method. This process was conducted with a 10x Genomics Visium CytAssist workflow using clinical-grade H&E staining on a Leica Bond instrument, followed by coverslipping and 40x WSI on the Aperio GT450 platform. Spatial transcriptomics data were filtered to include the top 1000 most variable genes across slides identified by SpatialDE ^14^. Separately, 708 WSIs were collected from colorectal cancer patients from the Dartmouth Hitchcock Medical Center, for whom, histological stage annotations were available. Finally, WSIs were obtained for a cohort of 350 colorectal cancer patients from The Cancer Genome Atlas (TCGA) for whom survival information and lymph node metastasis information was available. All WSIs were stain normalized using the Macenko ^15^ method. Collected WSIs were split into non overlapping 224 × 224 patches via the PathflowAI Python package ^16^, whose embeddings served as node attributes in a graph. We compared several methods described below to encode information for these patches, which is the main focus of this study. Nodes were connected with edges based on spatial adjacency using the *knn_graph* (k-nearest neighbor) method from the torch_cluster Python package, with n=16. Patients from the in-house dataset and TCGA were separately partitioned into training, validation, and testing sets using a random 80/10/10 split. The collected datasets and the downstream tasks they were used on, are summarized below:

1. **Visium spatial transcriptomics slides (n=4; 20**,**000 spots/patches; Co-Registered Spatial Transcriptomics, H&E WSI):** to pretrain contrastive crossmodal model
2. **Dartmouth Hitchcock Medical Center (n=708 H&E WSI):** used for histological stage prediction and clustering analysis
3. **TCGA Cohort (n=350 H&E WSI):** used for lymph node metastasis prediction, survival prognostication, and tumor infiltrating lymphocyte (TIL) alignment analysis

### 2.2. Patch Level Pretraining Methods

Three embedding production methods were compared for the 224x224 patches used as nodes of the graphs representing WSI.

#### 2.2.1. ImageNet-Pretrained ResNet18

A ResNet18 CNN model pre trained on the ImageNet dataset (commonly used for embedding histopathology patches) was accessed using the *torchvision* Python package (https://github.com/pytorch/vision). The model was truncated through the penultimate layer, to extract length 512 vectors/embeddings for each input patch.

#### 2.2.2. Ciga Self Supervised Histopathology Pretrained ResNet18

A separate ResNet18 CNN model pretrained using a self-supervised learning (SSL) SimCLR ^17^ contrastive procedure on histpathological imaging datasets was similarly accessed and truncated through the penultimate layer to extract length 512 embeddings for all patches. In summary, SimCLR employs an objective function that encourages similarity between embeddings from augmented (i.e., “corrupted”) views of the same image, while penalizing based on dissimilarity between views from different images. This model was made publicly available by Ciga et al ^10^, and has been previously shown to outperform the aforementioned ImageNet-pretrained model on a variety of downstream modeling tasks.

#### 2.2.3. Spatial Omics-driven Crossmodal Pretrained Encoder

A contrastive cross-modal model encoding image patches and spatial transcriptomic profiles was created, similar to the model implemented by Zeng et al ^18^. Input images patches of size 224x224 were encoded into embeddings of size 512 units, using the feature extraction portion of a CNN initialized with weights from the ResNet model trained by Ciga et al. Spatial transcriptomics profiles containing expression of the most spatially variable 1000 genes across Visium slides, selected to avoid overfitting on genes with imprecise expression, were encoded with three standard fully connected (FC) layers of size 512. The embeddings from co-registered patches from each modality (ST, WSI) were passed through a common projection layer of size 512, to output a single embedding per modality (ie; one vector of length 512 which describes an image patch, and one of length 512 which describes the corresponding gene expression). Crossmodal and unimodal contrastive penalties are applied using the SimCLR loss function ^17^; during training, several augmentation strategies were applied to both the image patches and corresponding transcriptomic profiles to generate “corrupted” representations of each data type as means for comparison. Transcriptomic profiles were randomly masked and corrupted with noise with 30% probability. Images were augmented using a series of random flips, color jitter transforms, random grayscaling, random rotation, and random image solarization. Both the original and augmented image patches and transcriptomics profiles were encoded using the aforementioned neural network layers. The loss mechanism penalizes the model based on the difference between the embeddings from the original and augmented data from each modality. A crossmodal loss is used to maximize the similarity between the corrupted image and transcriptomic embeddings from the same patch. These three loss functions (augmented image to image, augmented transcriptomics to transcriptomics, augmented image to augmented transcriptomics) were summed to optimize the crossmodal contrastive model. This model was trained for 150 epochs with a batch size of 8 and a learning rate of 0.00001.

Visium sections corresponding to six patients were partitioned into the training set, and tissue sections from two patients were partitioned to the validation set. Validation set loss was used to inform selection of the top model, following training. The RELU activation function was applied to outputs of every layer. The image encoder pretrained using the co-registered transcriptomics information was retained for subsequent analysis, and was used to embed image patches which GNN models were to operate on. The usage of this image encoder derived using this training protocol for other ancillary tasks is the primary focus of this study. This model is further described along with data collection procedure, in Figure 1.

**Figure 1:**
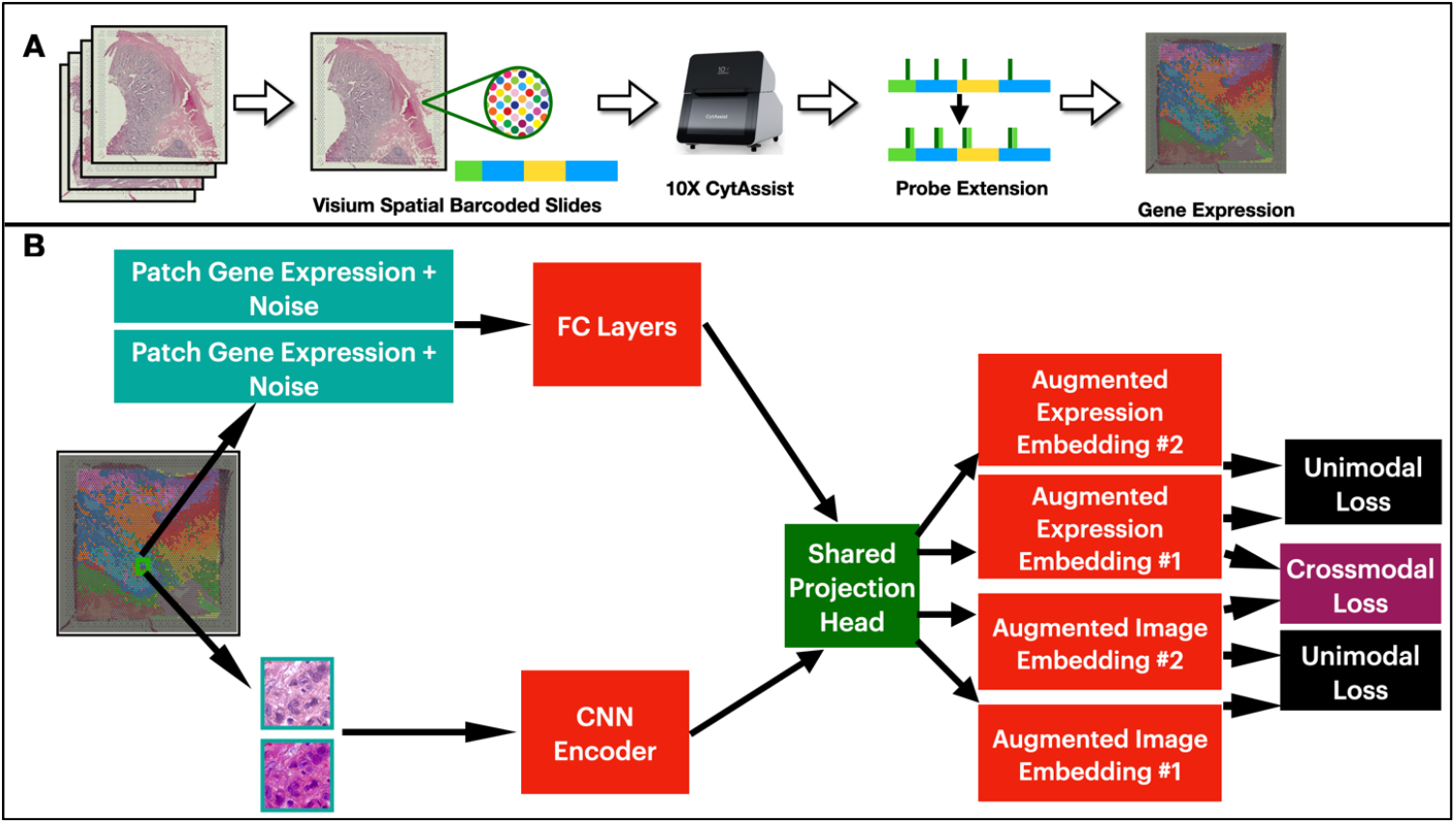
**A)** Data collection protocol for Visium spatial transcriptomics slide. **B)** Training protocol for spatial omics-driven crossmodal contrastive model; two views are generated per modality, per patch; each view is passed through the corresponding branch of the crossmodal model; embeddings are transformed using a shared projection head; unimodal and crossmodal contrastive losses are applied to output embeddings.

### 2.3. Downstream Outcome Prediction

We sought to understand whether CNN encoders, pretrained on co-registered spatial transcriptomics data, could enhance the predictions on a range of different GCN tasks. A graph convolutional network was constructed to take an input graph of nodes represented by length 512 embeddings, followed by three GCNConv graph convolutional layers ^19^ to contextualize and aggregate embeddings into length 128, with SAGEPooling pooling ^20^ layers (ie: 30% of patches retained, for subsequent layers; SAGEPooling stochastically samples higher-order neighborhoods of patches) placed after each convolutional layer. These pooling layers learn to downsample graphs, to push the model to learn focused information relevant to the training task. Graph embeddings were aggregated using global mean pooling after each SAGEPooling layer. These embeddings were combined using the JumpingKnowledge mechanism, resulting in a single vector of length 128 to represent the entire input graph/WSI. Finally, two fully connected layers were applied to this embedding, followed by a single output layer. The model (**Figure 2**) was applied to the following prognostication-focused experiments/outputs to assess patch encoding mechanisms:

**Figure 2:**
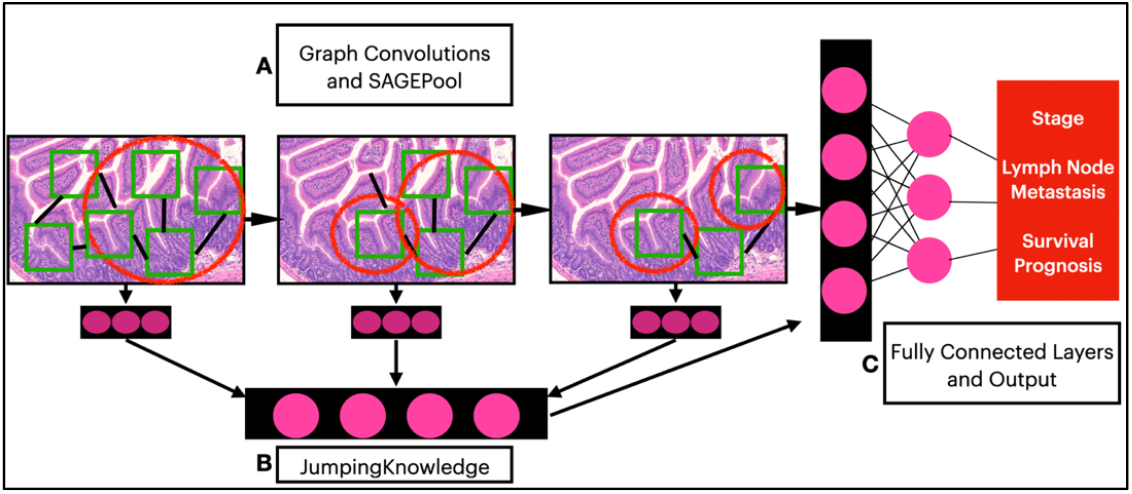
Overview of generalized GCN for downstream outcome modeling; initial patch embeddings vary across experimentation.

#### 2.3.1. Histological Stage Prediction

The in-house dataset was used to train and assess model capability to predict dichotomized tumor histological stage (T-stage; signifies depth of invasion) - low (stage 0, stage 1, stage 2) or high (stage 3, stage 4). A sigmoid function was applied to the output of the final layer in the GCN, and model training was supervised using a binary crossentropy loss function.

#### 2.3.2. Survival Prognostication

The TCGA dataset was used to train and evaluate GCNs to assess for time to death using hazard predictions, indicating the real-time risk of death. Model training was supervised using a standard Cox loss, which considers the predicted risk, patient censor status, and duration (either days to death or days to last follow up). This setup entails the proportional hazards assumption, that predictors have a constant hazard ratio (i.e., relative risk between two patient groups) over time.

All GCN models were trained for up to 30 epochs, using a learning rate of 0.001 and batch size 8. Top model checkpoints were selected for evaluation following training, based on validation set loss. GCN models were implemented using the Pytorch Geometric ^21^ Python package. Three separate GCN models were trained for each prediction task - one for each patch embedding mechanism. Stage prediction and lymph node metastasis models were evaluated on held-out test sets using F1-score and area under the curve (AUC), while C-index was used to evaluate prognostication models. These metrics are reported using 95% confidence interval derived from 1000 sample non-parametric bootstrapping procedures.

### 2.4. Embedding Clustering Quality Analysis

The ability of patch embeddings to capture morphological and molecular heterogeneity across slides was assessed across embedding methods, using an unsupervised clustering approach and the in-house dataset. For each WSI in the dataset, KMeans clustering (k=5; chosen via coarse optimization to ensure stability when run numerous times) was applied to the patch embeddings derived by each pretraining method (standard ResNet, Ciga et al, spatial pretrained) to elucidate sub-groups of patches implicitly captured by the representations. Clusters were plotted across slides to visually ensure that they represented different morphologies and structures within slides. Subsequently, the Calinski-Harabanz (CH) index ^22^ and the Davies-Bouldin (DB) index ^23^ were computed for the clustering result for each pretraining strategy. The ANOVA-based CH score assesses the density and separation of clusters, with a higher value indicating greater density within clusters and separation among different clusters. Similarly, the DB index measures the ratio between within-cluster and cross-cluster separation. Thus, superior patch embeddings should result in a relatively high CH index and low DB index. The per-WSI scores were used to calculate average CH index and DB score at a 95% confidence interval, for each pretraining method.

### 2.5. TIL-based Model Interpretation

Previous research has demonstrated the importance of tumor infiltrating lymphocytes (TILs) and the tumor microenvironment on the progression of colon cancer ^24^. We sought to demonstrate the interpretability of GCN models developed here using the TCGA dataset, by comparing regions of WSI given high attention with previously published predicted TIL maps ^25^ for corresponding slides. Patches deemed important by GCN models trained on lymph node metastasis prediction were determined by extracting patches remaining in WSI graphs following the final pooling layer; for a given patch, being left in its graph by a GCN model following three pooling layers, indicates its significance to the model. The coordinates of these patches were compared to those describing the locations of predicted TILs via Wald Wolfowitz testing ^26^, where the null hypothesis would indicate high overlap between these two sets of coordinates. Accordingly, Wald Wolfowitz testing was used to calculate a test statistic per slide per GCN model trained with each patch embedding method– negative values of this test statistics, *W*, represents the localization of TILs. Spearman’s rank correlation coefficients (alpha p-value = 0.05) were calculated to evaluate the relationship between the test statistic (*W*), and predicted hazard. A negative correlation coefficient would suggest a statistically significant association between predicted hazard and TIL spatial localization, following biological knowledge holding that TILs help inhibit colon cancer proliferation and migration ^27^. Test statistics were further dichotomized to indicate presence/lack of TIL localization, to compare these relationships across the GCN model using embeddings derived from the Ciga et al method, versus the model using spatially pretrained embeddings.

## 3. Results^†^

### 3.1. Quantitative Predictive Analysis

Held out testing-set performance for GCNs trained to predict stage, lymph node metastasis, and survival prognosis, are presented in **Table 1**; models which used patch embeddings derived from the spatial omics-driven mechanism outperformed those using the compared methods for all three experiments.

**Table 1:** Test set performance metrics (95% confidence interval) of GCNs trained using various patch embedding mechanisms, for binary stage prediction, lymph node metastasis prediction, and survival prognostication.

| Task | Measure | ImageNet ResNet | Ciga et al ResNet | Spatial Pretrained |
| --- | --- | --- | --- | --- |
| Stage Prediction | AUC | 0.935 ± 0.003 | 0.948 ± 0.002 | **0.981 ± 0.001** |
|  | F1-Score | 0.863 ± 0.004 | 0.858 ± 0.004 | **0.878 ± 0.004** |
| Lymph Node Metastasis | AUC | 0.651 ± 0.004 | 0.612 ± 0.004 | **0.708 ± 0.003** |
|  | F1-Score | 0.560 ± 0.002 | 0.630 ± 0.003 | **0.671 ± 0.005** |
| Survival Prognostication | C-index | 0.597 ± 0.003 | 0.582 ± 0.002 | **0.638 ± 0.002** |

For the classification experiments, models using embeddings derived from the spatial omics-driven mechanism outperformed those which used embeddings from the ImageNet-trained ResNet18 CNN by an average of 6.98% measured by AUC, and outperformed models using embeddings derived from the ResNet18 pretrained by Ciga et al, by average of 9.47%. GCNs using spatial omics-driven embeddings (C-index 0.638) also outperformed ImageNet-trained ResNet18 embeddings (C-index 0.597) and embeddings derived from the model trained by Ciga et al (C-index 0.582).

### 3.2. Clustering Evaluation

A KMeans clustering approach paired with CH index and DB index calculation was employed to compare the abilities of these different pretraining approaches to elucidate molecular and morphological heterogeneity across slides; the results of this analysis are presented in **Table 2**. An example visualization including regions of a slide assigned to clusters indicating by different coloring, is presented in **Figure 3**; additional examples are available in Supplementary Figures S2 and S3.

**Table 2:** Clustering quality metrics calculated across embedding methods.

| Measure | ImageNet ResNet | Ciga et al ResNet | Spatial Pretrained |
| --- | --- | --- | --- |
| Calinski-Harabanz Index | 643.76 ± 17.51 | 786.70 ± 20.40 | **2605.68 ± 70.66** |
| Davies-Bouldin Index | 1.90 ± 0.01 | 1.719 ± 0.01 | **0.975 ± 0.01** |

**Figure 3:**
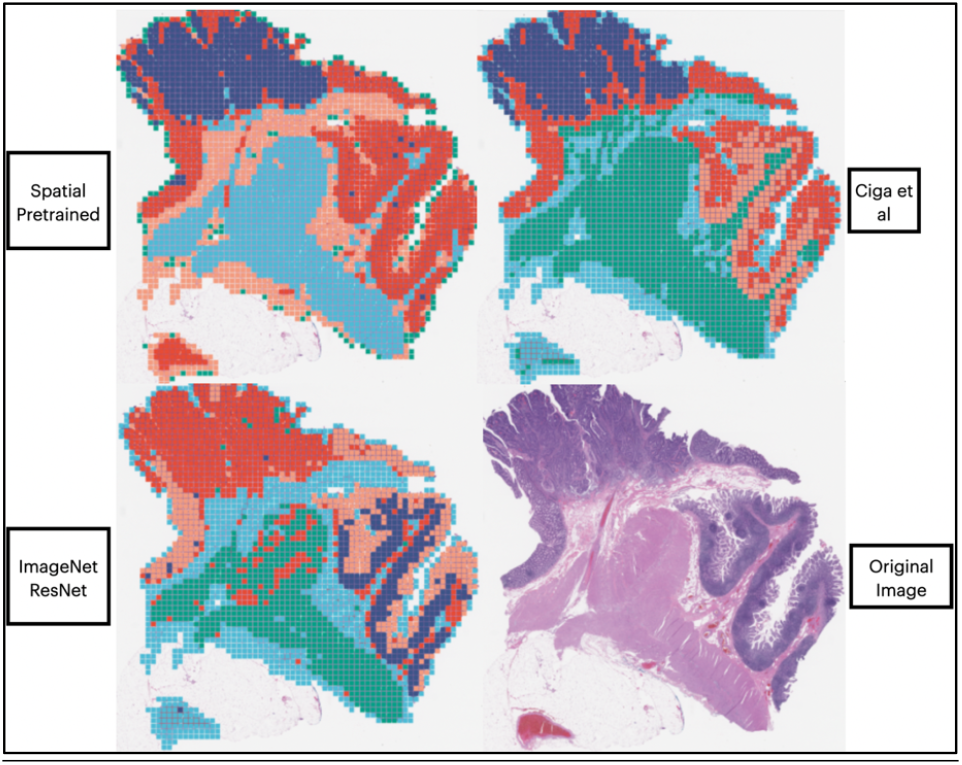
Example visualization of clustering of embeddings derived using various methods, for a single WSI.

Embeddings from the contrastive crossmodal spatial model resulted in a significantly higher CH index and lower DB index, versus both the ImageNet-pretrained ResNet and the ResNet trained on histopathology datasets via self-supervised learning by Ciga et al.

### 3.3. Model Interpretation

Spearman’s correlation coefficient values testing the relationship between lymph node metastasis risk predicted by GCN models using various patch embedding mechanisms and TIL localization elucidated via Wald Wolfowitz testing, are presented in **Table 3** along with corresponding p-values, suggesting both the Ciga and spatial pretrained models were able to derive TIL-associated embeddings related to instantaneous hazards. Boxplot visualizations comparison of predicted model risk and dichotomized TIL alignment are presented in **Supplementary Figure S4**.

**Table 3:** Spearman’s correlation coefficient values for TIL localization versus predicted lymph node metastasis risk, across GCN models using various patch embedding methods.

|  | ImageNet ResNet | Ciga et al ResNet | Spatial Pretrained |
| --- | --- | --- | --- |
| **Spearman’s Coefficient** | -0.061 | -0.426 | -0.218 |
| **Spearman’s P-value** | 0.2693 | 2.2e-16 | 7.74e-5 |

## 4. Discussion and Conclusion

This is the first study which aims to determine whether leveraging spatial omics data to pretrain image patch encoders using a cross modal contrastive mechanism can improve downstream performance in graph convolutional networks, which may improve automated cancer patient analysis. We compared spatial omics-driven embeddings against those extracted from a standard ResNet18 CNN pretrained on the ImageNet dataset, and a ResNet18 pretrained using self-supervised learning on histopathology datasets. GCN models trained and evaluated using the spatially enhanced embeddings outperformed those using the baseline embedding methods on three downstream tasks – stage prediction, lymph node metastasis prediction, and prognostication. This suggests that incorporating spatial transcriptomics information into the pretraining process of image patch encoders, enhances the quality of learned representations, beyond what is extracted from state-of-the-art techniques which use solely images for patch encoding pretraining.

Additional quantitative analysis from clustering patch embeddings indicates that the models leveraging spatially-pretrained embeddings were superior at capturing distinct heterogeneities across slides, versus models using patch embeddings from existing strategies. Thus, we expect future applications of the developed spatial pretraining method for patch embeddings, to improve the performance of workflows aiming to capture tissue heterogeneity, including tumor subcompartment segmentation.

Furthermore, Wald Wolfowitz testing paired with Spearman’s correlation coefficients, suggests that GCN models using embeddings from the spatial pretraining method and the Ciga et al method, learned to highlight TILs to contextualize prognostic assessment of cancerous tissue when considering lymph node metastatic potential, particularly in patients whom the models understood to be at lower risk. The Spearman’s coefficient value for the GCN model using ImageNet ResNet patch representations was markedly closer to 0 versus the other two methods, indicating far weaker correlation in this relationship. Interestingly, the magnitude of the coefficient for the GCN model using the Ciga et al embeddings was nearly double that of the spatially pretrained embeddings, indicating that the Ciga et al method may induce greater tendency to turn to TILs for understanding patient profiles. That such nuances can be extrapolated, demonstrates the interpretability of graph-based modeling for cancer histopathology, and further emphasizes the importance of enhancing the ability of such methods.

Overall, our results indicate that spatial omics data can be effectively mined in a crossmodal fashion, to improve existing image-based deep learning workflows to analyze cancer histopathology; this also adds to the growing body of literature ^28–30^ which reflects the importance of enhancing pretraining mechanisms as a basis of improving deep learning models for cancer histopathology.

A key limitation of this study is the relatively small dataset used to pretrain the spatially-enhanced crossmodal contrastive model; spatial transcriptomics data was only generated for 8 total slides due to high resource and time costs and the limited size of the tissue placement area on Visium slides. Furthermore, coarse hyperparameter search was used to select GCN architecture parameters, as a detailed experiment here was beyond the scope of this study. Future works will seek to use larger cohorts to pretrain the spatial model to improve quality of extracted embeddings. Additionally, the embeddings from the spatially enhanced model can evaluated for use in applications other than GCNs, such as Transformer networks – which have become popular in cancer histopathology in recent years ^31,32^ – histology image search, and multimodal data integration.

## 5. Acknowledgements and Location of Supplementary Material

The results published here are in part based on data generated by the TCGA Research Network: https://cancer.gov/tcga. The authors acknowledge the support of the Center for Clinical Genomics and Advanced Technology in the Department of Pathology and Laboratory Medicine of the Dartmouth Hitchcock Health System which includes the Pathology Shared Resource, at the Dartmouth Cancer Center with NCI Cancer Center Support Grant 5P30 CA023108-37. Spatial transcriptomics assays were carried out in the Genomics and Molecular Biology Shared Resource (GMBSR) at Dartmouth which is supported by NCI Cancer Center Support Grant 5P30CA023108 and NIH S10 (1S10OD030242) awards. Spatial studies were conducted through the Dartmouth Center for Quantitative Biology in collaboration with the GMBSR with support from NIGMS (P20GM130454) and NIH S10 (S10OD025235) awards. Supplementary materials can be found at the following DOI: https://doi.org/10.5281/zenodo.8197573.

## Supplementary Material

**Table S1:** Hyperparameters for models trained during this study.

|  | Contrastive Crossmodal Spatial Pretraining | Downstream Outcome GCNs (n=3) |
| --- | --- | --- |
| Epochs | 150 | 30 |
| Learning Rate | 0.00001 | 0.001 |
| Batch Size | 8 | 8 |

**Figure S2:**
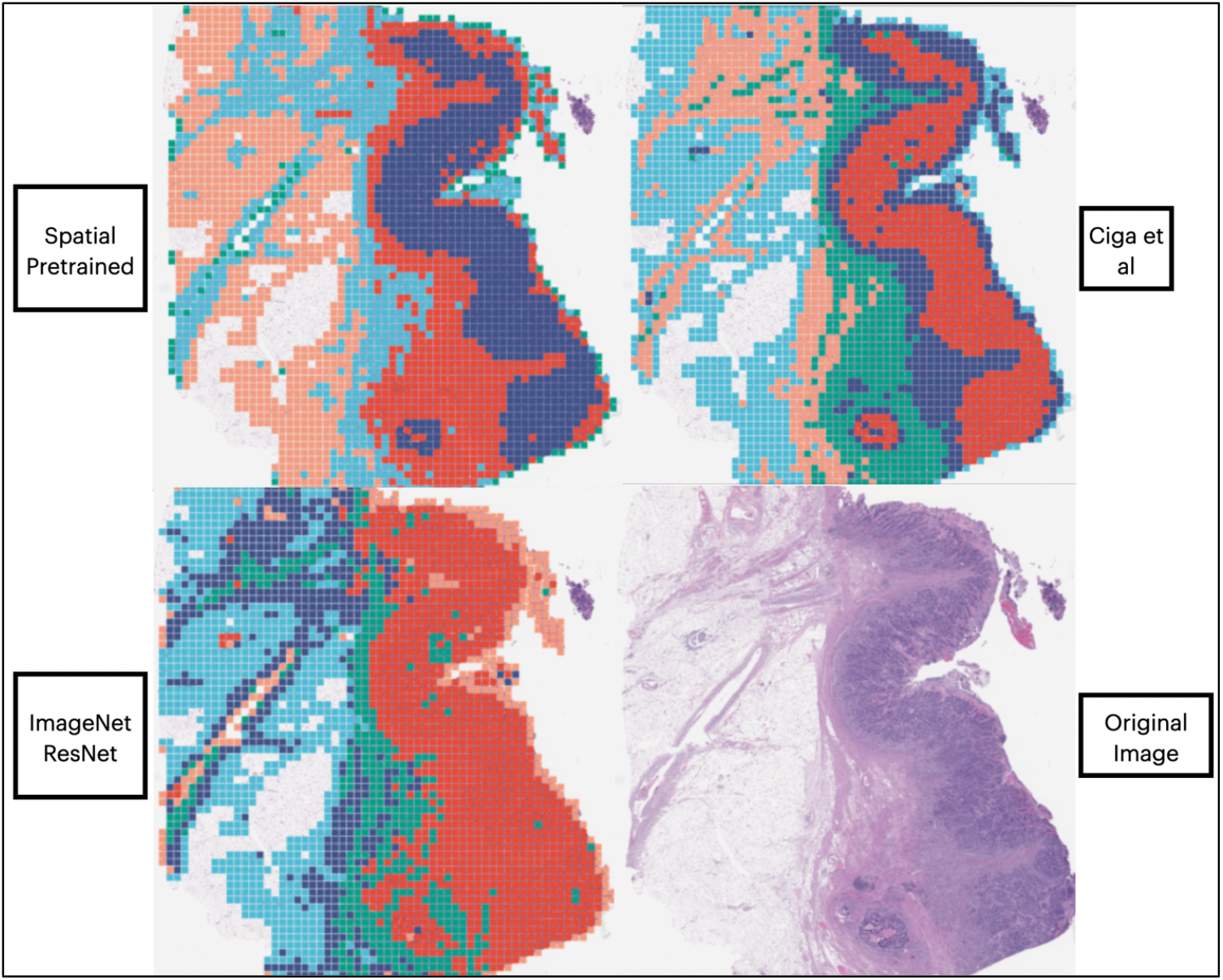
Additional example visualizations of KMeans clustering-derived subgroups in WSI, identified using various patch embedding mechanisms.

**Figure S3:**
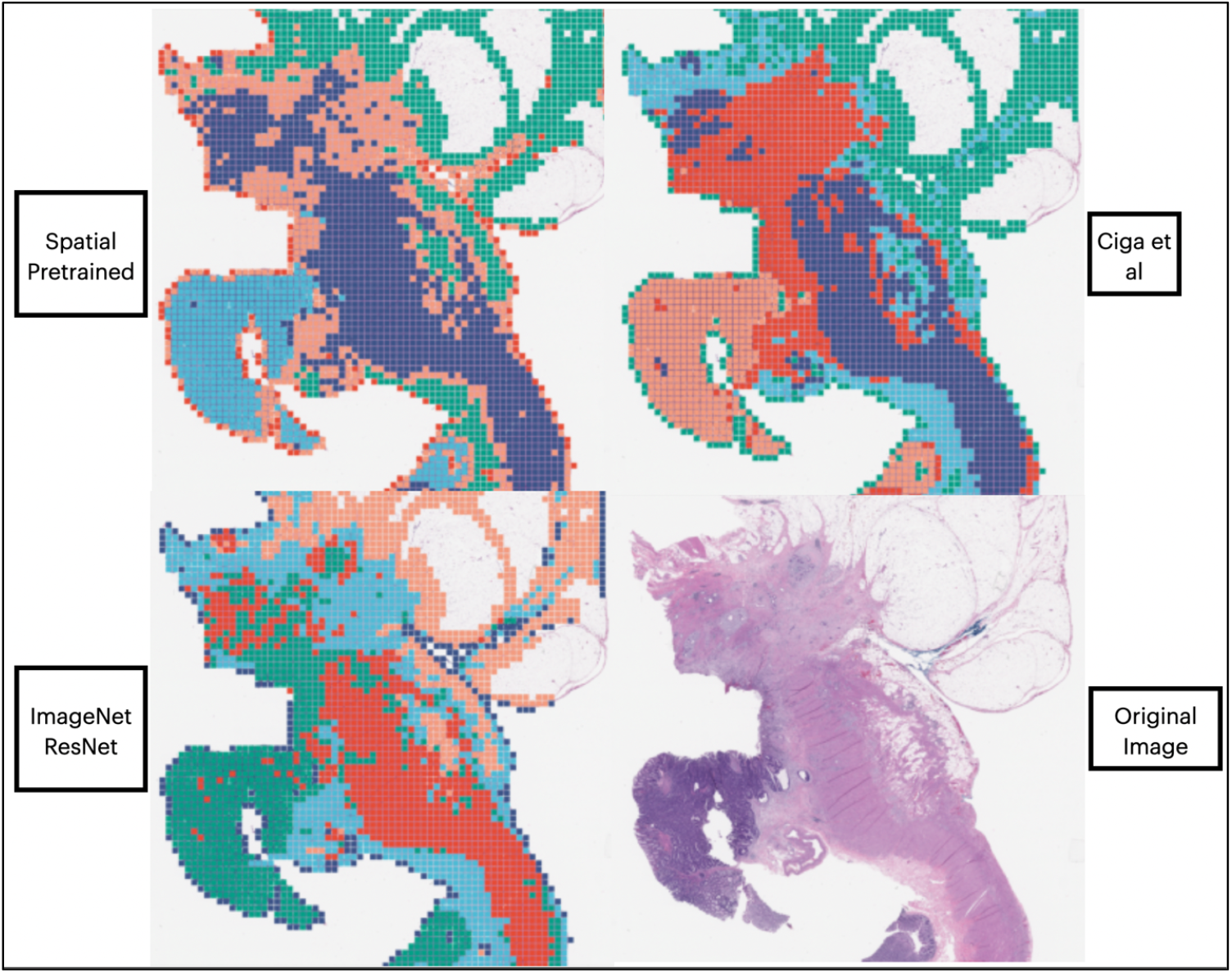
Additional example visualizations of KMeans clustering-derived subgroups in WSI, identified using various patch embedding mechanisms.

**Figure S4:**
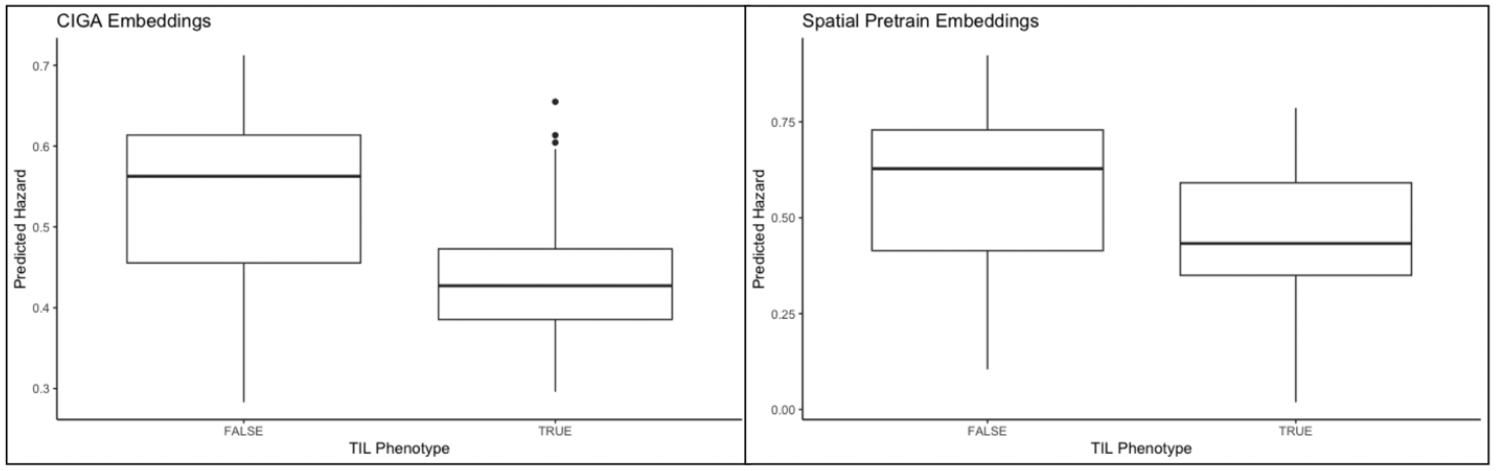
Boxplot visualizations of predicted lymph node metastasis risk versus localization status of tumor infiltrating lymphocytes (TILs) by GCN models utilizing patch embeddings derived by the Ciga et al method, and the developed spatial crossmodal pretraining method.

## Footnotes

† Supplementary materials can be found at the following DOI: https://doi.org/10.5281/zenodo.8197573.

